# Interaction of hypoxia and nicotine acetylcholine receptor signaling network reveals a novel mechanism for lung adenocarcinoma progression in never-smokers

**DOI:** 10.1101/2021.09.08.459287

**Authors:** Namita Pandey, Jonita Chongtham, Soumyadip Pal, Anant Mohan, Tapasya Srivastava

**Author notes:** **Corresponding author:** Tapasya Srivastava; Room No. 202, Department of Genetics, University of Delhi South Campus, New Delhi – 110021, 91-9810777253. **Emails of all authors:** NP; JC, SP, AM; TS. **Author Contributions:** NP: Concept and design, methodology and experiments, analysis, writing manuscript; JC: experiments and validation; SP: Analysis; AM: Data acquisition, storage and analysis; TS: Concept and design, resource and funding, writing manuscript and editing, correspondence.

## Abstract

High incident of lung cancer among never smokers and their disease pathogenesis is an unexplained phenomenon. We have analyzed 1727 lung cancer patient data to understand the impact of smoking on overall survival of lung cancer patients and have observed a difference of only 47 days between smokers and never smokers in adenocarcinoma patients suggesting that the disease is equally fatal in never-smokers irrespective of gender. In this study, we have investigated the possible collaboration between the nAChR and hypoxia signaling pathway to elucidate a mechanism of disease progression in never-smokers. We report a previously unidentified increase in both acetylcholine and *nAChR-α7* levels in non small cell lung cancer cells in hypoxia. Similar increase in ubiquitously expressed *nAChR-α7* transcripts was also observed in other cancer lines. A direct binding of HIF-1α with the hypoxia response element (HRE) present at -48 position preceding the transcriptional start site in *nAChR-α7* promoter region was established. Significantly, the increased acetylcholine levels in hypoxia drove a feedback loop via modulation of PI3K/AKT pathway to stabilize HIF-1α in hypoxia. Further, Bungarotoxin, an antagonist of nAChR-α7 significantly reversed hypoxia mediated metastasis and induction of HIF-1α in these cells. Our study gives a plausible explanation for the equally worse prognosis of lung adenocarcinoma in never-smokers wherein the nAChR signaling is enhanced in hypoxia by acetylcholine, in the absence of nicotine.

## 1. Introduction

A baffling and unanswered gap in our understanding of lung cancer progression is the development of the disease in never-smokers. In lung adenocarcinoma, these patients were observed to have a more aggressive form of tumor at the time of diagnosis [1], that metastasized more frequently to the lung pleura [2]. Globally, as well as in India, there has been a considerable increase in the number of lung cancer - especially adenocarcinoma - cases in non-smokers [3]. The growing number of lung cancer cases in never-smokers, their unique molecular profile and response to therapy prompted us to study non-tobacco related risk factors and disease progression in never-smokers [4].

nAChRs are ubiquitously expressed [5] ligand-gated cation channels that form pentameric structures assembled from a family of subunits that include α1-α10 and β1-β4 encoded by the CHRNA gene family [6]. Importantly, they are activated in response to both endogenous neurotransmitter acetylcholine (ACh) and exogenous agonists, such as nicotine or tobacco-specific nitrosamines NNN (N’-Nitrosonornicotine or NNK (4-(methylnitrosamino)-1-(3-pyridyl)-1-butanone)[7]. nAChRs in lung cancer cells act as central mediators for various signaling pathway like PI3K/AKT mediated cell survival pathway, Raf-MAPK mediated cell proliferation pathway, ERK dependent calpains phosphorylation pathway leading to increased invasion and migration and stimulation of VEGF expression causing tumor angiogenesis [8-10].

This work proposes a possible explanation for the undetermined lung adenocarcinoma disease progression in never-smokers by scrutinizing the nAchR pathway and HIF signaling pathways. Their individual contribution to worsening disease prognosis is well known [11,12], yet their association has not been previously determined.

## 2. Materials and Methods

### 2.1. Patient data collection, follow up and relevant clinical characteristics

Patients visiting the Lung Cancer Clinic of the Department of Pulmonary Medicine from 2011 till 2018 have been included in the analysis [14]. Data entry of 1727 patients for whom the date of confirmed diagnosis/registration/treatment, definitive date of death/last follow-up and smoking status was available.

### 2.2. Cell culture

The human non-small cell lung cancer (NSCLC) cell lines A549 (adenocarcinoma), NCI-H460 (NSCLC), CHO (mammalian epithelial line), A172 (glioma) and AGS (gastric cancer) were purchased from National Centre for Cell Science (NCCS, Pune, India) and cultured in Dulbecco’s Modified Eagle Medium (DMEM) (Gibco, Thermo Fisher Scientific, USA) supplemented with 10% Fetal Bovine Serum (FBS) (Gibco, Thermo Fisher Scientific, USA) and 5µg/ml ciprofloxacin (Sigma, USA) at 37°C in a humidified incubator with 5% CO_2._ The *in vitro* 0.2% hypoxic conditions were created as reported before (13). Various treatment conditions such as Nicotine (Sigma, US), α-Bungarotoxin (αBTX) (Sigma, US) and Acetylcholine chloride (Ach) (Invitrogen, US) in doses as indicated in the text were given in normoxia or 0.2% hypoxia.

### 2.3. Acetylcholine receptor activity assay

Acetylcholine concentration in cell lysate was detected using Amplex® Red Acetylcholine/ Acetylcholinesterase (AChE) Assay Kit (Invitrogen, US) using the manufacturer’s protocol. Primer sequences used for cloning in Dual Luciferase Assay:

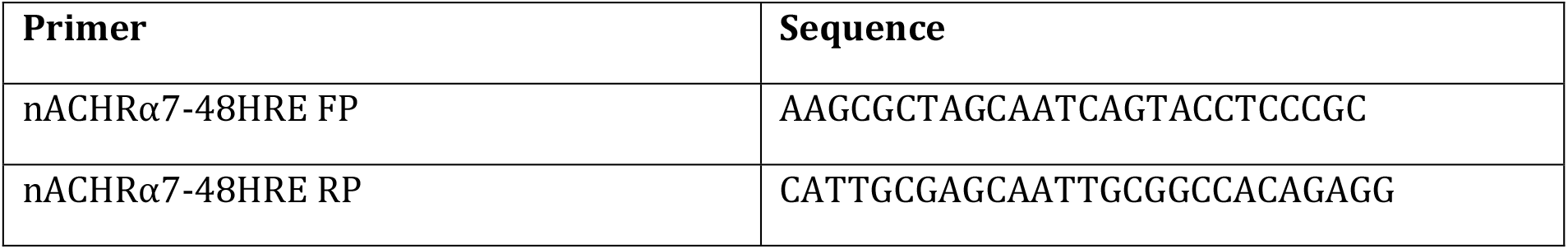

### 2.4. Immunofluorescence for detection of nAChR expression

A549 cells seeded in four well chamber slides were grown in normoxia and 0.2% hypoxia for 4hr at 37°C. Cells were then quickly washed with PBS, fixed in 4% paraformaldehyde for 10min and blocked in 3% BSA for 2hrs. Cells were then incubated with rabbit-anti-nAChR- α7 antibody, Alexa Fluor 594 (Invitrogen, USA) (1:100) and stained with 4,6-diamidino-2-phenylindole (DAPI) (Sigma, USA). The slides were mounted with Fluoroshield™ (Sigma, US), sealed, and observed under the fluorescence microscope (Olympus, USA).

### 2.5. Chromatin immunoprecipitation (ChIP) assay

A549 cells cultured in normoxia and hypoxia for 16 hours were fixed with one-tenth volume of crosslinking buffer and incubated at room temperature for 10 minutes. Immunoprecipitation was done with 4μg of HIF-1α (BD Biosciences) and HIF-2α (Cell Signaling Technology) antibodies in the presence of magnetic DynaBeads (Invitrogen) at 4°C overnight. Amplification of different segments of the regulatory regions of nAChR-α7 was done by PCR using specific primers. The PCR conditions include an initial cycle of 95°C for 5 min, followed by 35 cycles of 30s at 95°C, 30 sec at annealing temperature and 30s at 72°C with a final extension step of 72°C for 10 minutes. Reactions were normalized with Input DNA while pull down using anti-IgG antibody served as negative control.

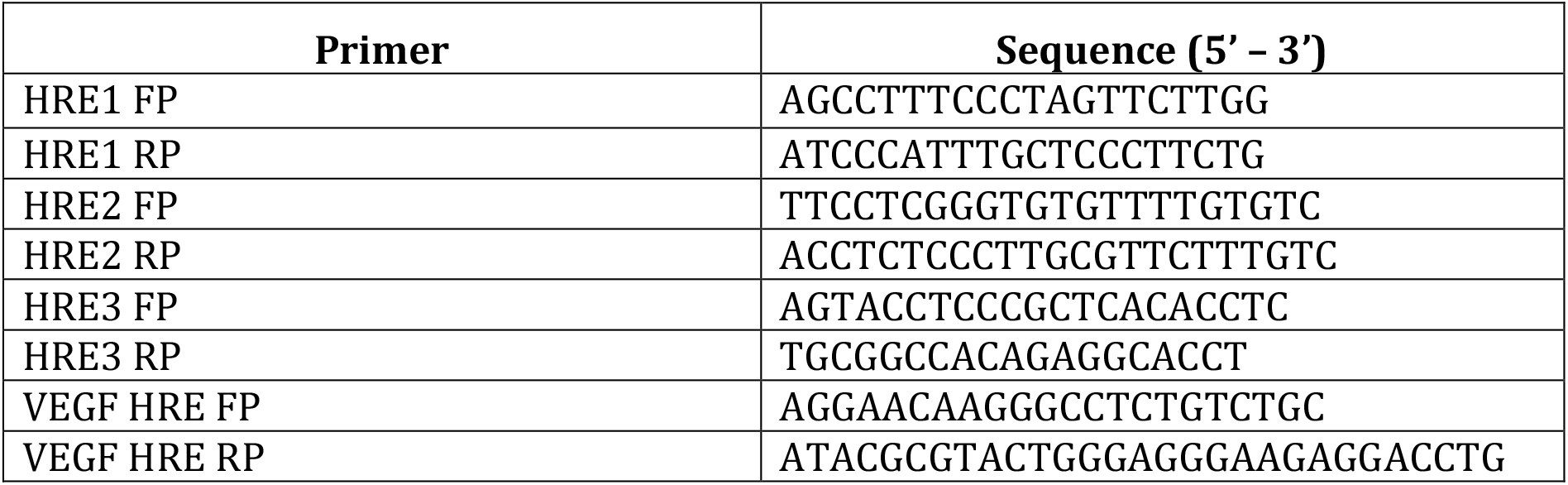

### 2.6. Transient Overexpression of HIF1α

Cells grown in 6 well plate was transfected with HIF1α mutant plasmid (HA-HIF1αP402A/P564A/N803ApcDNA3) using Lipofectamine LTX (ThermoScientific, US) according to manufacturer’s instructions. This plasmid could continuously express HIF1α containing P402A/P564A/N803A mutation and therefore will not undergo rapid degradation even in normoxia [15].

### 2.7. Dual Luciferase Assay

The proximal HRE containing region of nAChR-α7 was PCR amplified and cloned into pGL3basic vector in *NheI* and *XhoI* sites. pGL3-nAChRα7-48HRE was then co-transfected with pGL4-basic (10:1 ratio) in A549 cells seeded in a 24-well plate and incubated in 0.2% hypoxia for 48 hours. Cell lysates were prepared, and luciferase activity was measured according to manufacturer’s protocol (Dual Luciferase Reporter Assay System, Promega) using a microplate reader (Tecan Infinite 200 Pro).

### 2.8. Wound healing and migration assay

For wound heal (scratch) assay, cells were seeded in 24-well plate at density of 1×10^5^ cells per well. The images were captured at 0hr and 24hr to identify the migration rate of treated cells compared to the control.

In transwell migration assay, about 200µl of serum free media containing 5X 10^4^ cells were placed on the upper chamber of a transwell insert. Transwell inserts were washed in PBS several times to remove excess stain and filter membrane was carefully cleansed with cotton swab. The images were captured at a magnification of 20X in different fields of view and number of cells migrated through the membrane were counted using the ImageJ software.

### 2.9. RNA isolation and Real-time PCR

RNA was extracted using Trizol (Invitrogen, US) according to the manufacturer’s instructions. About 2μg of total RNA was reverse transcribed using SuperScript III reverse transcriptase (Invitrogen, US) and quantitative PCR were performed in Rotor-Gene Q (Qiagen, Germany). The primer sequences are given below:

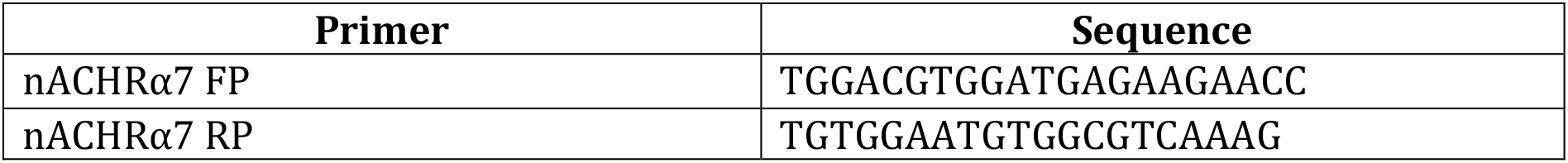

### 2.10. Western Blotting analysis for protein expression

Cells were washed with PBS and homogenized using lysis buffer. Protein estimation, SDS-PAGE and electrotransfer were performed using standard protocol. Immunoblots were detected with the chemi-luminescence reagent Luminol (Thermo Scientific, US), and developed by ImageQuant LAS4000 (Fujifilm, Japan). β-Actin was used as loading control for the total protein content. Antibodies used in the study are given below.

CHRNA7 – ab216485 (Abcam, US)

β-actin – NB600-501 (Novus Biologicals, US)

HIF1α – 610958 (BD Biosciences, US)

HIF2α – 7096 (CST, US)

pAKT – 4060 (CST, US)

### 2.11. Statistical analysis

All experiments were performed in at least three biological replicates for reproducibility. Bars in the graph represent +standard deviation of at least three independent experiments. Two-tailed Student t test was used to analyze the difference between two samples and p < 0.05 was considered as statistically significant. ANOVA was performed on patient data to understand if the difference in overall survival (OS) arises from smoking history. This has been calculated for males and females in different histological groups. Analysis of variance compares variance within a group i.e smokers and non-smokers separately versus variance between the two groups. F-test has been performed and p value has been calculated as significant if < 0.05.

## 3. Results

### 3.1. Association of late-stage lung cancer progression to smoking status and insignificant difference in overall survival of smokers and never-smokers in adenocarcinoma

From the AIIMS cohort, 1727 patient data was evaluated for a correlation of smoking status in different histological types in those patients for whom first diagnosis/date of registration and date of death/last follow-up was available. The number of never-smokers in histological sub-types squamous cell carcinoma (SqCC), non small cell carcinoma (NSCC) and small cell carcinoma (SCLC) were much fewer compared to those in adenocarcinoma (AC). OS in smokers (186 days) was significantly lower than in non-smokers (232 days); p = 0.0005 [F = 12; F_crit_ = 4.9] in cumulative histological cancers analyzed. OS in smokers and non-smokers of lung adenocarcinoma with a sizable number of 246 non-smokers and 333 smokers was analyzed for overall survival (OS) separately. OS of lung adenocarcinoma never-smokers (258 days) was higher than that of smokers (211 days); p= 0.02 (F = 4.9; F_crit_ = 3.85). Although the p-value is < 0.05, the limited 47 days difference suggests that disease is equally fatal in never-smoking adenocarcinoma patients and points at involvement of genetic and environmental factors, other than smoking, in the disease progression (Figure 1B). The difference in OS between smokers and never-smokers in males and females was not significantly different in any of the groups, and the never-smoker females with adenocarcinoma did not have an exceptional advantage in disease progression (Figure 1C). The mechanism of disease progression in smokers and non-smokers appear to be equally fatal.

**Figure 1:**
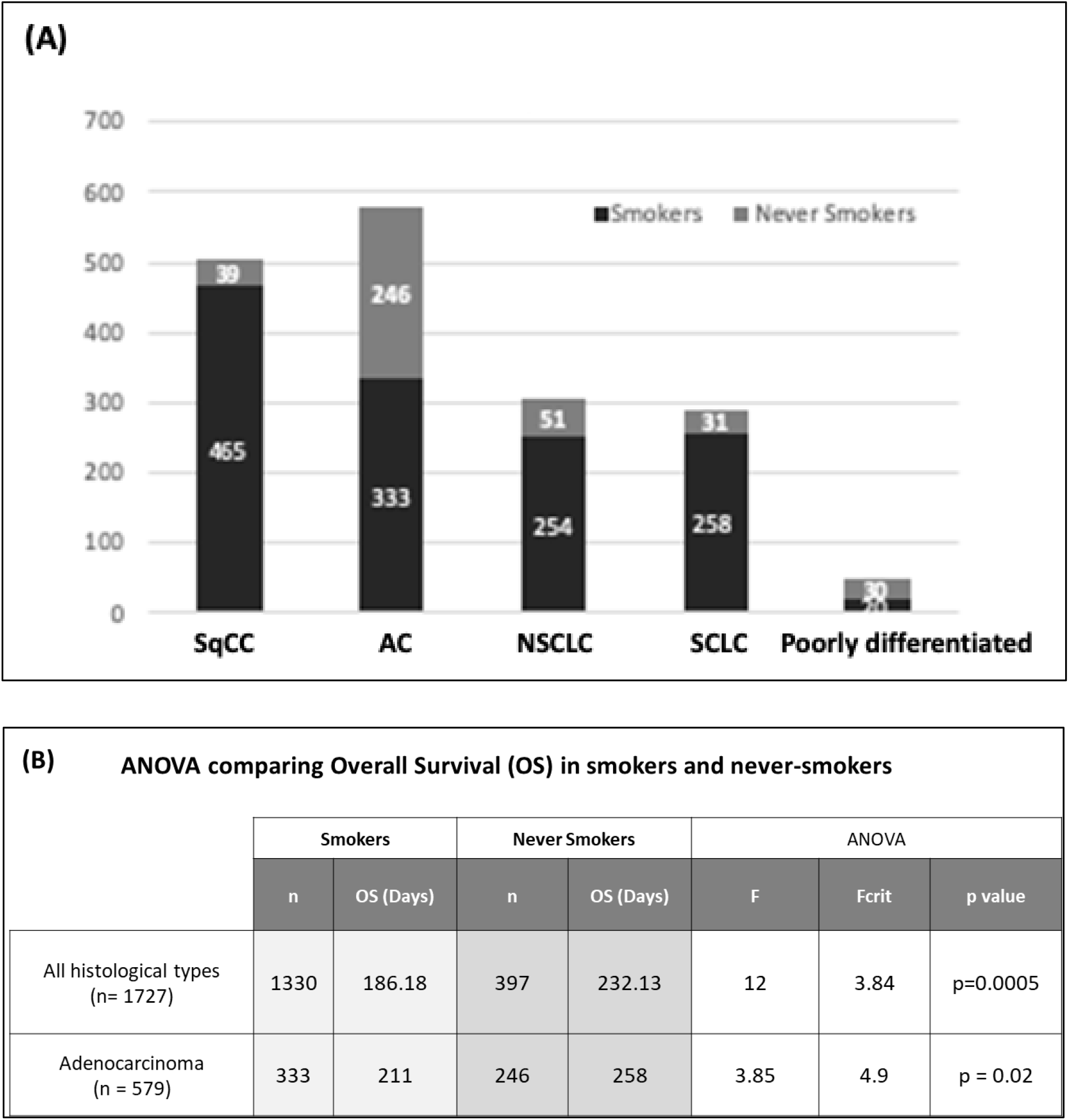
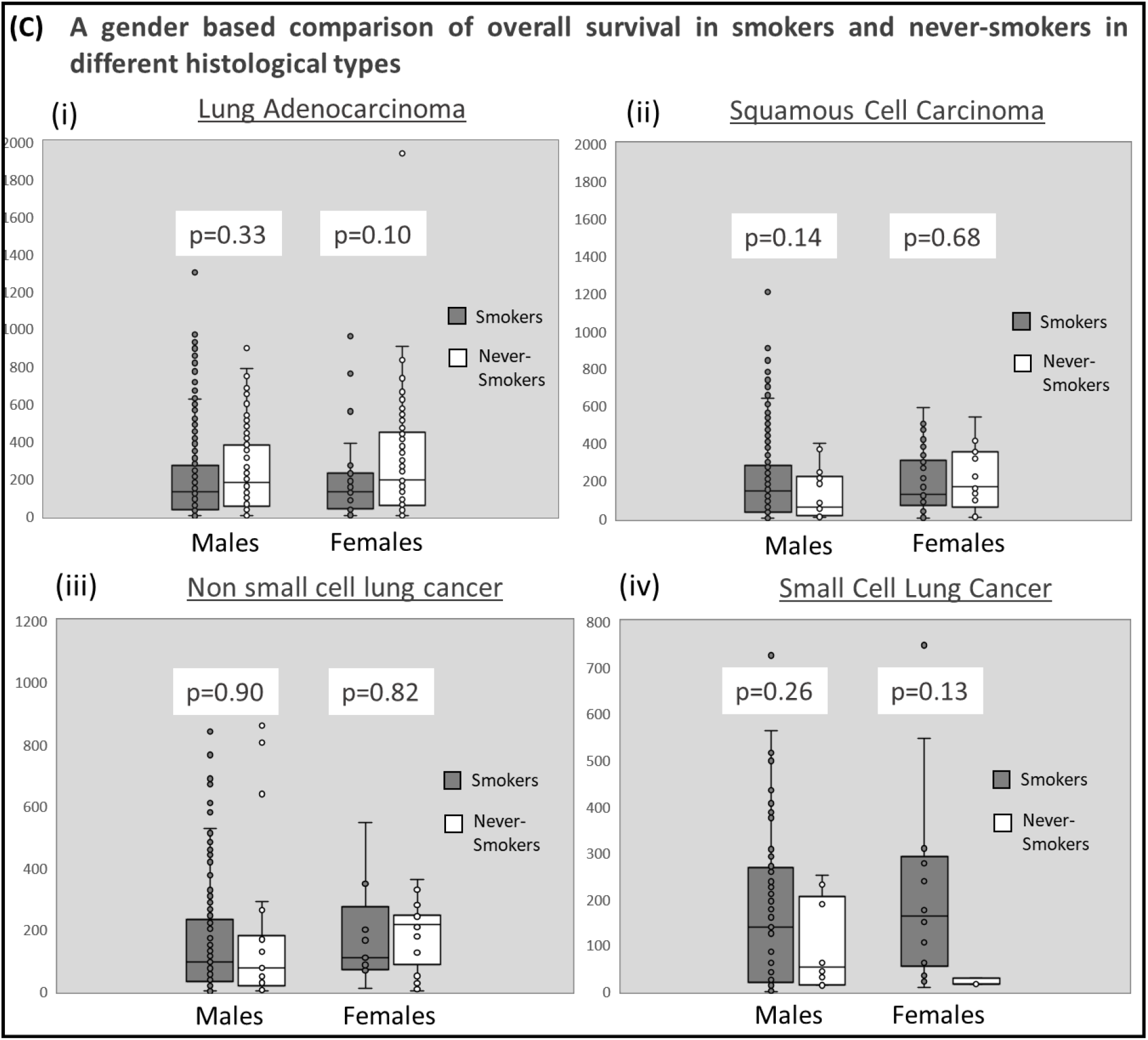
Analysis of smoking status, gender and overall survival: **(A)** Distribution of smoking status in different histological types analyzed in 1727 records of patients from 2011 to 2018. SqCC: Squamous cell carcinoma; AC: Adenocarcinoma; NSCLC: Non small cell lung carcinoma; SCLC: small cell lung carcinoma **(B)** ANOVA comparing Overall Survival (OS) for never-smokers and smokers for all histological types and adenocarcinoma. **(C)** ANOVA comparing overall survival in different histological types reveals no difference in overall survival between never-smokers and smokers among both males and females. Particularly in the case of lung adenocarcinoma (i) where a substantial number of never-smokers are females. In case of (iv) the never-smokers among females is a very small number and therefore, inconclusive

### 3.2. Direct binding by HIF-1α, and not HIF-2α, drives promoter activity and expression of *nAChR α7* in different cancer lines

A 1500bp promoter region upstream of the transcription start site of all nine *nAChR* genes was analyzed for the presence of HIF recognized hypoxia response element (HRE) and ancillary sequences. Several putative core HRE sites were identified [Figure 2A(i) and 2A(ii)]. However, the occurrence of both HRE consensus motifs (A/GCGTG) as well as ancillary sequences (CAGC) recognized by HIFs was observed in only *nAChR α7* gene at -48 position preceding the transcriptional start site of and was further pursued. Three potential HRE consensus motifs at positions -48, -389 and -1010 within the ∼1500-bp region preceding the transcriptional start site [Figure 2B] were investigated by chromatin immunoprecipitation. Cells exposed to 0.2% hypoxia showed binding of HIF-1α to the HRE consensus motifs at -48 position [Figure 2C(i)] but not to HRE site at -389 [Figure 2C(ii)] and -1010 position [Figure 2C(iii)]. We determined that HIF-1α, and not HIF-2α, specifically binds to and positively regulates *nAChR-α7* HRE consensus motifs at -48 position on its promoter region in hypoxia. (Figure 2C).

**Figure 2:**
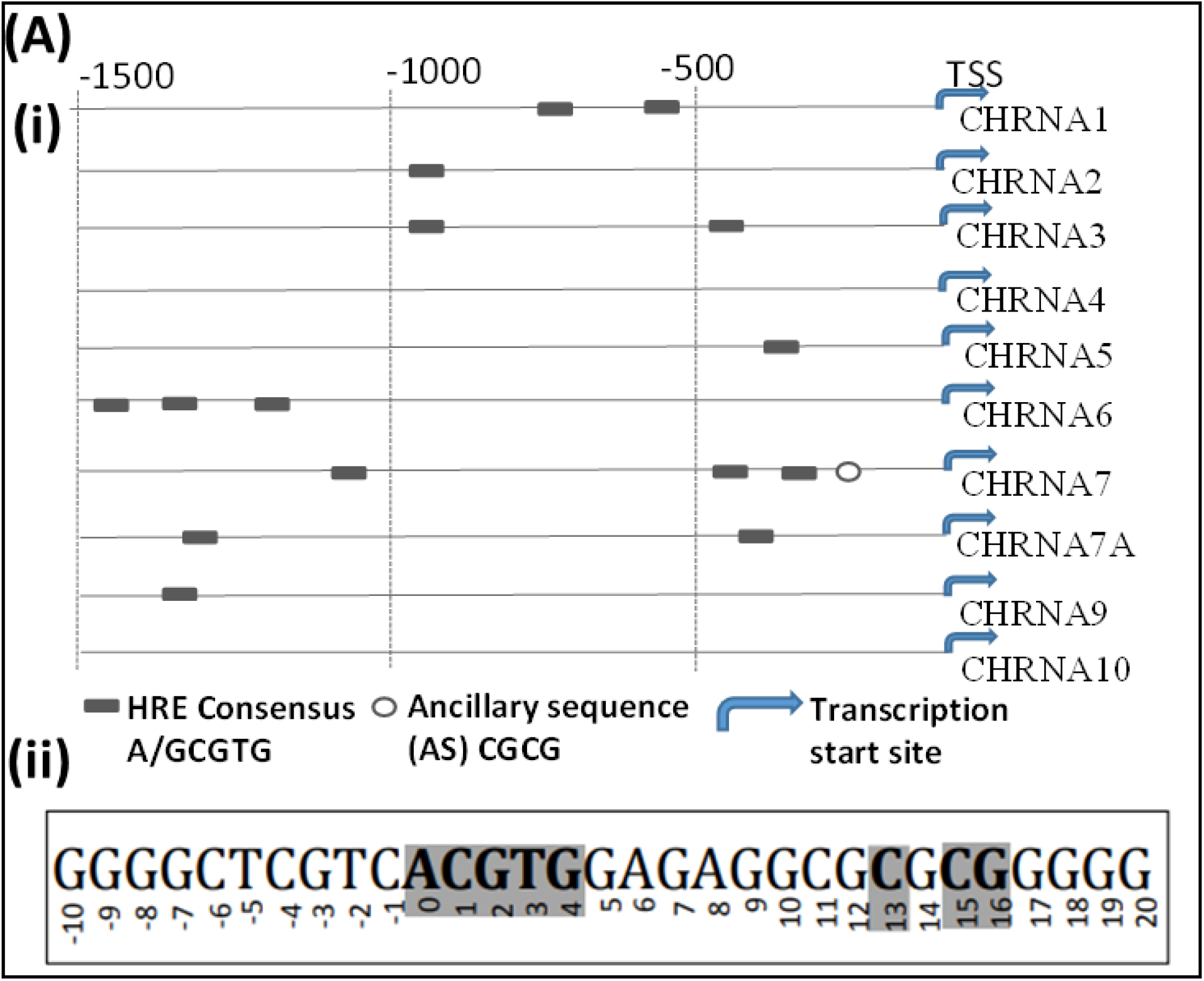
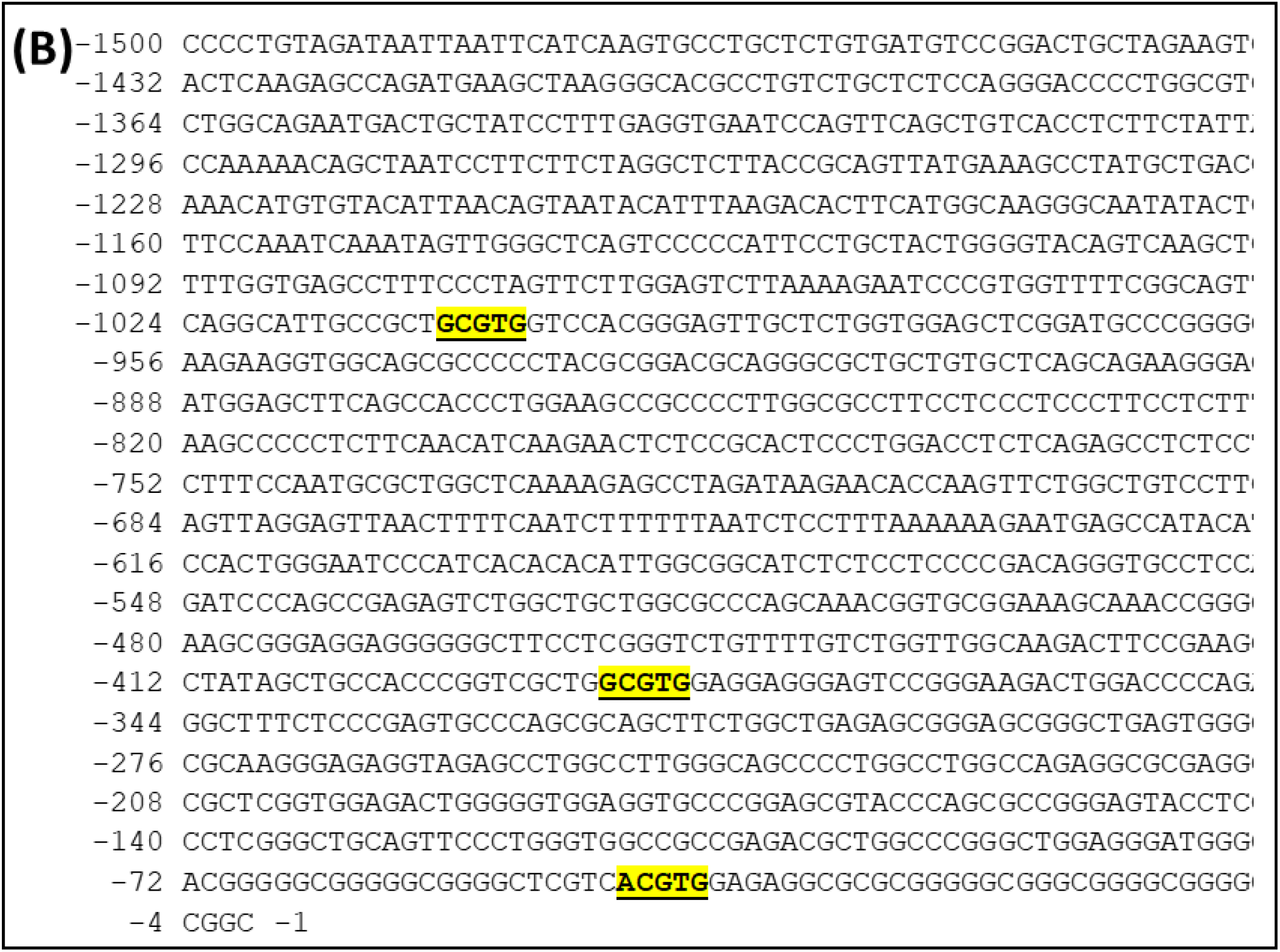
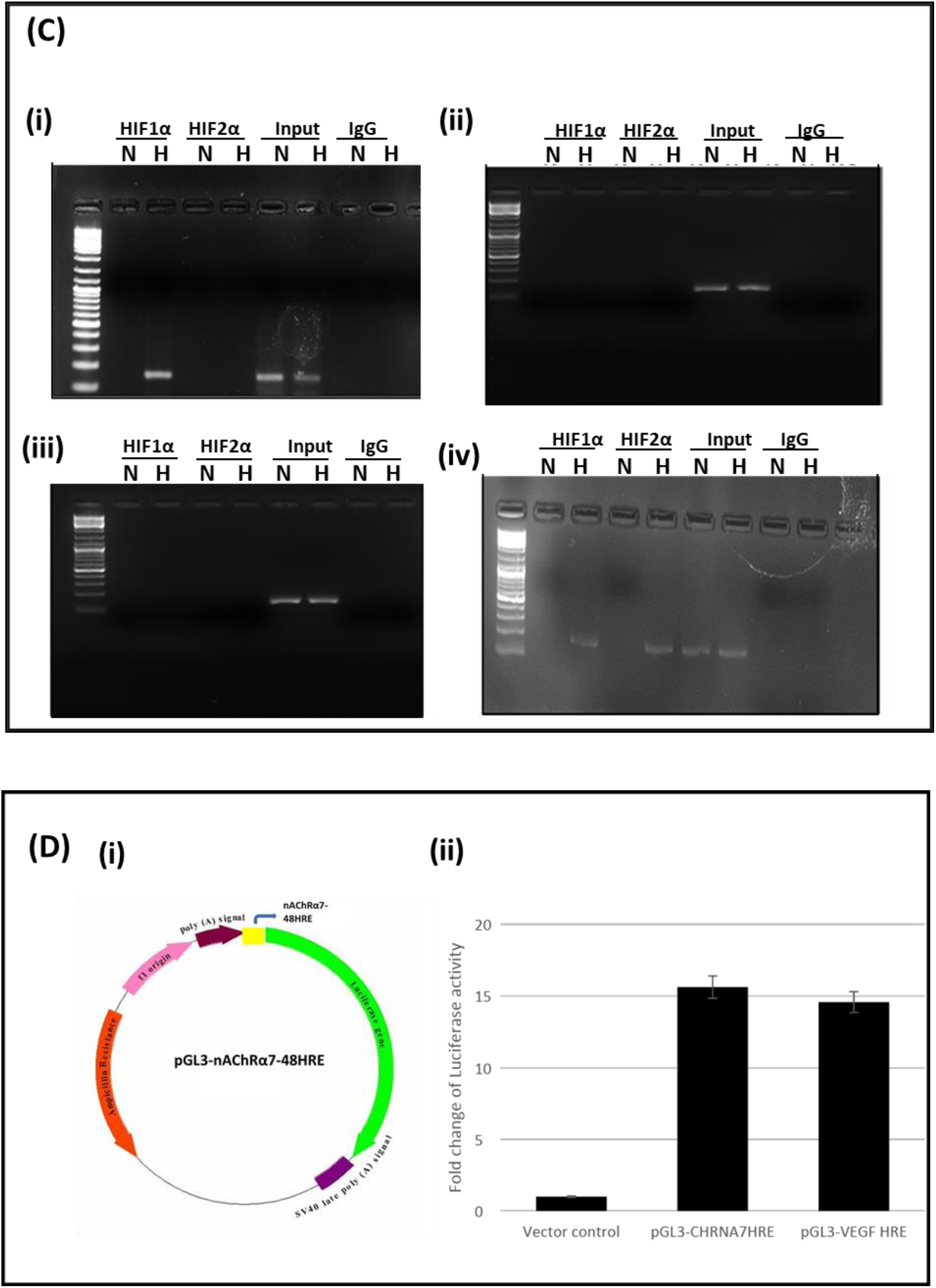
Identification of HRE sites and HIF-1α driven promoter activity in A549 lung cancer cells in hypoxia: **(A)** (i)Schematic diagram representing promoter region of nine CHRNA genes acquired from NCBI and analyzed manually for HRE and ancillary sequences recognized by HIFs. (ii) The potential HRE site at -48 position in the CHRNA7 promoter region possess preferred A base at position 0 and contain CGCG motif between +13 to +16 positions like the preferred CACG motif recognized by HIFs at this position **(B)** The nAChR-α7 promoter region containing three potential HRE site at -48, -389 and -1010 positions highlighted in yellow. **(C)** ChIP assay for HIF-1α and HIF-2α binding at the three potential HRE sites. HRE site at -48 position (i) was identified in hypoxia cultured cells but not at -389 position (ii) and - 1010 position (iii). Immunoprecipitation with HIF-2α antibody showed no enrichment of nAChR-α7 promoter fragment. Promoter fragment of VEGF (iv) with HRE was used as a positive control for HIF-1α/HIF-2α pull down. **(D)** Enhanced promoter activity was observed with nAChR-α7 genomic region containing putative HRE at -48 position (∼16 folds) as compared to vector control (ii). HRE region of VEGF was used as the positive control.

Similar to nicotine, hypoxia significantly enhances nAChR-α7 expression (Figure 3A, 3B). HIF-1α expressing A549 cells exhibit significant {10 fold (p<0.05)} increase in nAChR-α7 mRNA level similar to VEGF, a positively HIF-1α-regulated gene that increased ∼12 folds (p< 0.05) compared to the untransfected cells (Figure 3C). Besides A549, an increased expression of *nAChR-α7* was observed in another NSCLC line NCI-H460, mammalian epithelial CHO, A172 glioma and AGS gastric cancer cell lines (Figure 3D) in hypoxia.

**Figure 3:**
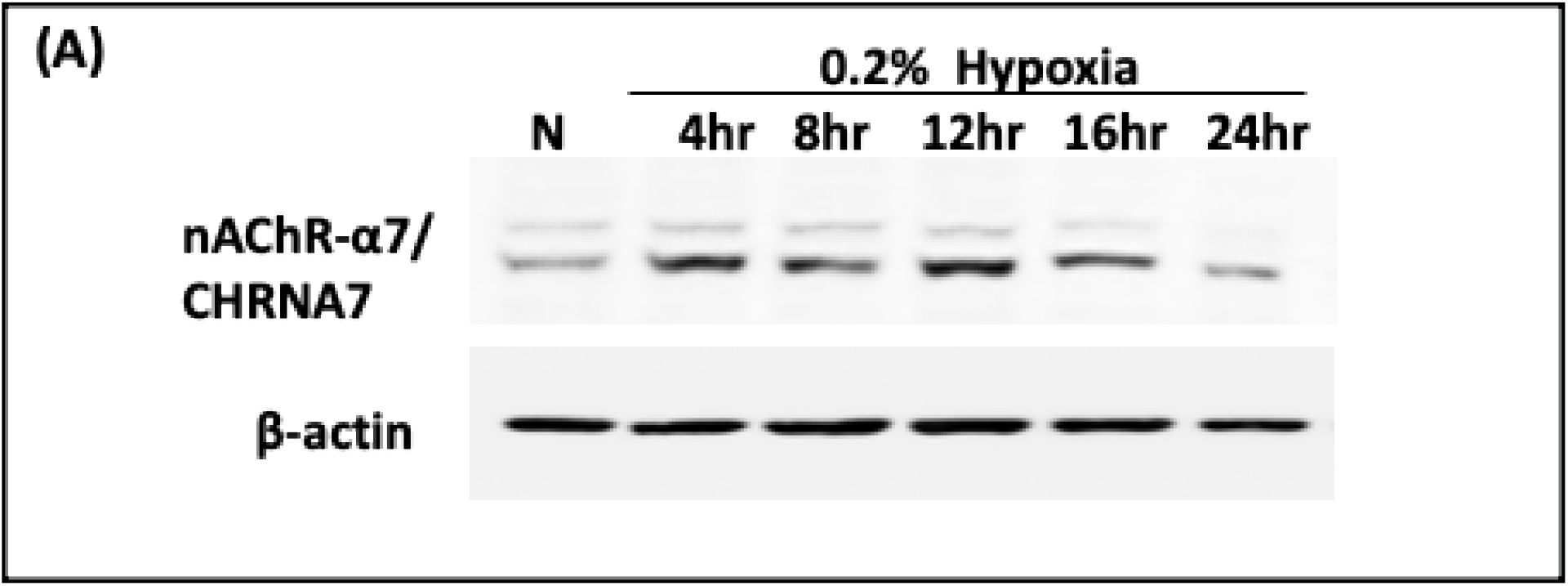
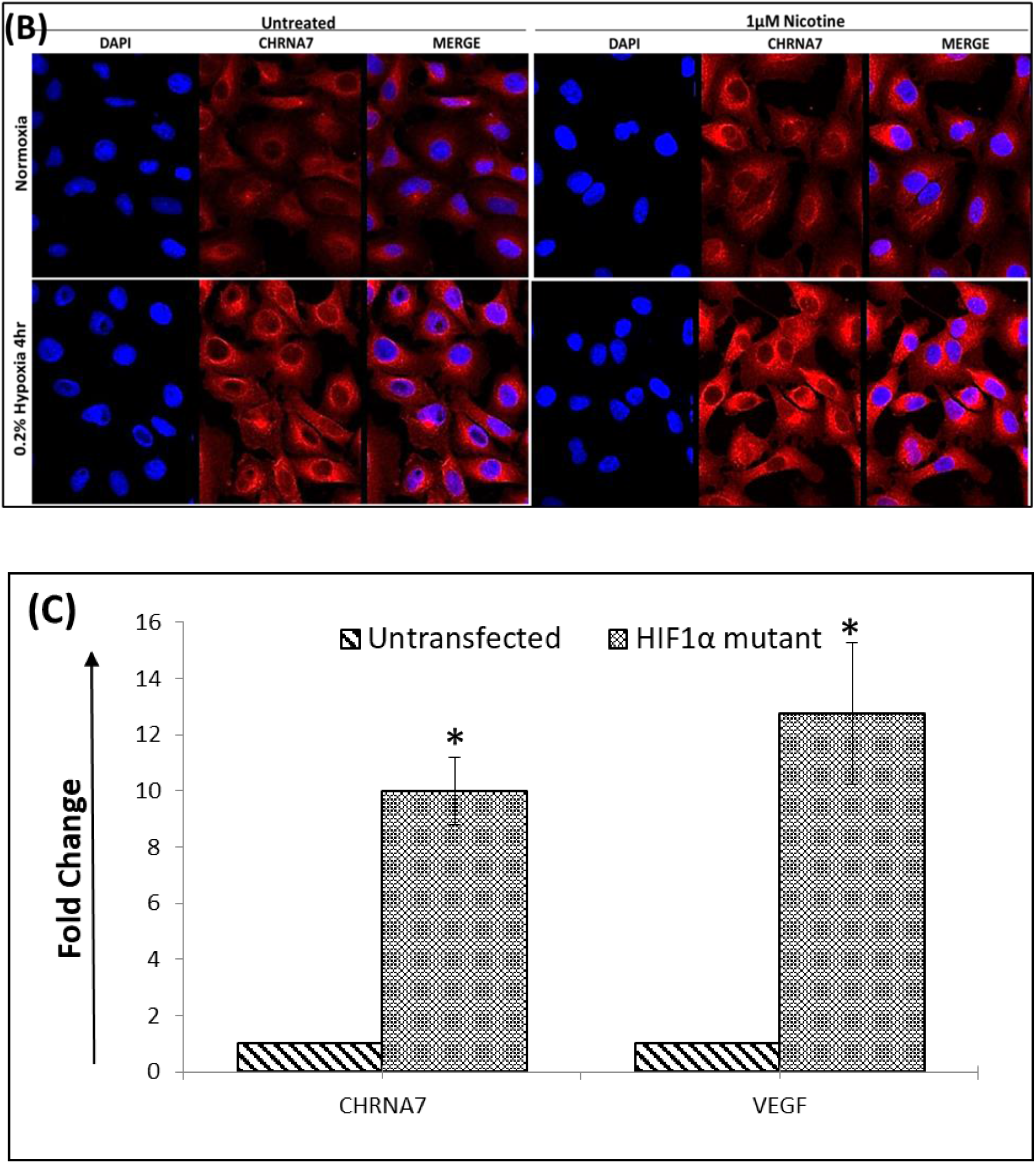
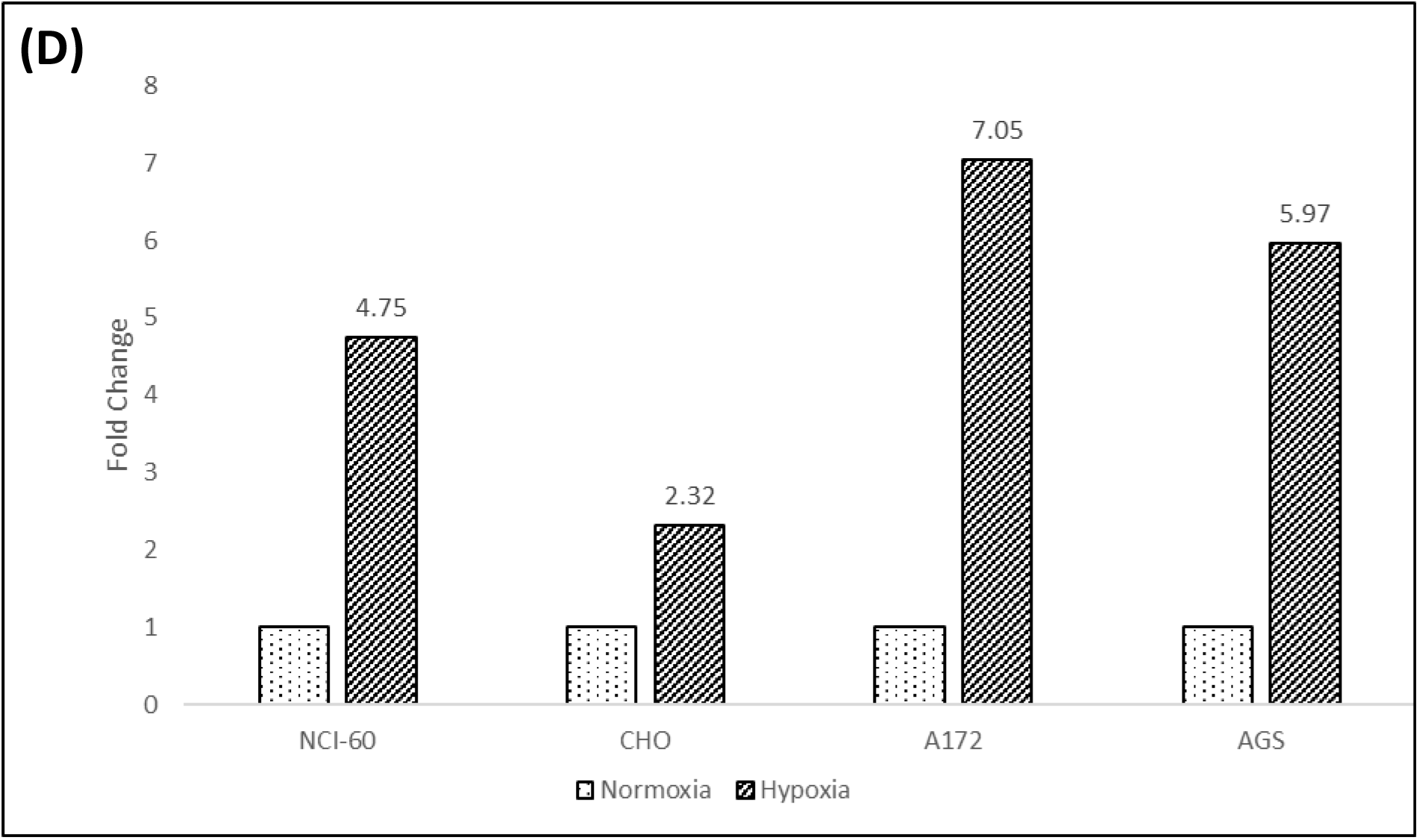
HIF-1α driven nAChR expression in A549 lung cancer cells in hypoxia: **(A)** nAChR-α7 protein levels were elevated inA549 cells cultured in 0.2% hypoxia at different time points compared to normoxia (N). β–actin is used as loading control. **(B)** Immunofluorescence studies showing overexpressed nAChR-α7 protein in A549 cells in 0.2% hypoxia 4hr. Nicotine which is known to induce the mRNA and protein levels of nAChR-α7 was used as positive control. **(C)** HIF-1α overexpressing plasmids showed significant enhancement of nAChR-α7 transcripts similar to that of VEGF (positive control) **(D)** nAChR-α7 transcripts were enriched in NCI-H460, CHO, A172, and AGS cell lines in 0.2% hypoxia compared to normoxia (N).

### 3.3. nAChR-α7 mediated pathway regulates HIF-1α expression via induction of acetylcholine and enhances hypoxia induced migration

Acetylcholine, an endogenous ligand of nAChR-α7, is known to act as an autocrine and paracrine growth factor activating a feedback loop and stimulating cancer cell proliferation [7], and we investigated its association with the hypoxia signaling network.

We have identified a novel effect of hypoxia on acetylcholine production in lung cancer cells where acetylcholine was significantly (p<0.05) accumulated (from 4.5μM to 9μM) in 0.2% hypoxia in a time dependent manner (Figure 4A). Interestingly, *nAChR-α7* mediated signaling pathway reciprocates by increasing HIF-1α expression forming a feedback loop in hypoxic lung cancer cells. Significantly, we also show that acetylcholine, a ligand for *nAChR* significantly increased the expression of the stabilized HIF-1α in hypoxia in a dose dependent manner independent of nicotine (Fig 4B(i)). We found marked decrease in the expression of HIF-1α, but not HIF-2α in presence of *nAChR-α7* antagonist α-BTX {Figure 4B(ii)}. A noticeable increase in HIF-1α protein expression was observed in presence of *nAChR-α7* exogenous agonist, nicotine, in A549 lung cancer cells incubated in hypoxic condition {Figure 4B (iii)}. We observed a notable decrease of AKT phosphorylation with 25nM α-BTX treatment {Figure 4B(iii)}, at which the maximal reduction of HIF-1α protein level was also observed suggesting that in hypoxia *nAChR-α7* mediates accumulation of HIF-1α protein through PI3K/AKT pathway. nAChR mediated pathway contributes to hypoxia induced malignant phenotype and 25nM α-BTX significantly (p< 0.05) represses migration of A549 cells in hypoxia (Figure 4C, 4D), suggesting that *nAChR-α7* plays an important part in regulating hypoxia induced migration in A549 cancer cells.

**Figure 4:**
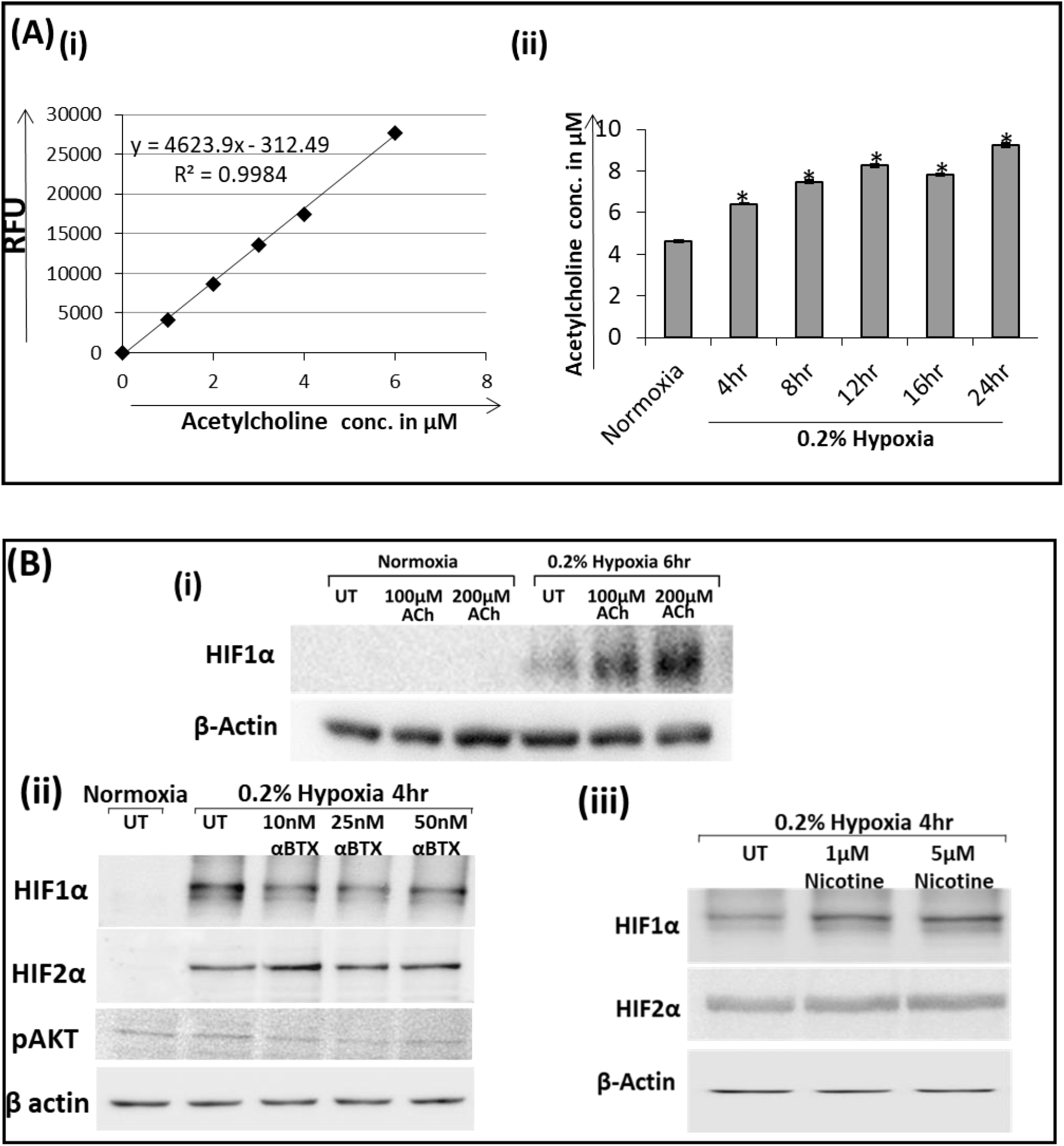
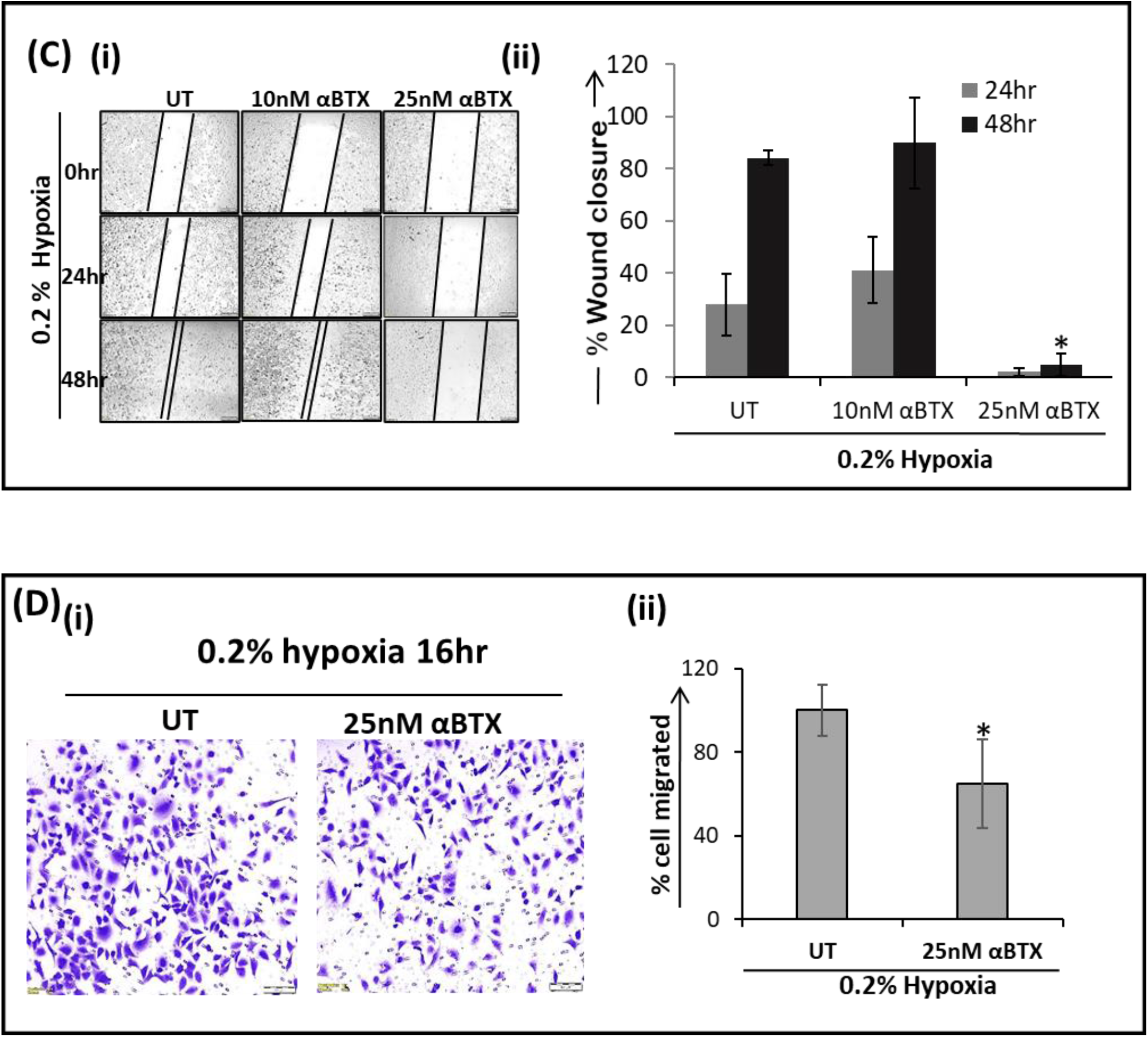
Role of nAChR-α7 mediated pathway in regulating the expression of HIFs and migration in A549 cancer cells in hypoxia: **(A)** Acetylcholine detection assay. (i) Acetylcholine standard curve (ii) Time point analysis of Acetylcholine concentration in normoxia and 0.2% hypoxia **(B)** Protein expression of (i) HIF-1α was increased with Acetylcholine (i) HIF-1α was increased and HIF-2α remained unchanged in presence of nAChR-α7 agonist nicotine (UT= untreated). (ii) HIF-1α and pAKT protein levels are repressed in presence of α-BTX. β–actin is used as loading control. **(C)** Wound healing assay. (i) Photomicrograph of wounded A549 cells treated with 10nM and 25nM α-BTX in hypoxic condition at different timepoints. (ii) Effect of α-BTX plotted as percentage of wound closure against time (0hr). **(D)** Migration assay. (i) Photomicrograph of migrated A549 cells 16hr after treatment with 25nM α-BTX in hypoxic condition (ii) Plot representing percentage of A549 cell migrated in untreated and α-BTX treated conditions. The error bars indicate standard deviations. * represents p< 0.05 relative to its untreated (UT) counterpart.

## 4. Discussion

The growing number of lung cancer cases in never smokers needs investigation of non-tobacco related risk factors and better understanding of the molecular mechanism of the disease [14,16]. Nicotine acetylcholine receptor activation with nicotine present in tobacco or endogenous ligand acetylcholine opens up downstream signaling associated with lung cancer risk and drug resistance. Elevated expression of nAChRs in lung cancer cells exemplifies their role in lung cancer progression. Various studies suggest that nAChR subunits (CHRNA1/*nAChR-α1*, CHRNA5/*nAChR-α5* and CHRNA7/*nAChR-α7*) are associated with a poor prognosis of lung cancer (17-19).

Our study establishes *nAChR-α7* as a novel target of HIF-1α transcription factor in lung cancer cells. It also successfully demonstrates that HIF-1α can regulate the *nAChR-α7* gene expression in lung cancer cells by directly binding to its promoter in hypoxia.

It has been suggested that the neurotransmitter, Acetylcholine functions as a growth factor in lung tumor progression by autocrine and/or paracrine signaling [20]. We observe a previously unreported acetylcholine accumulation in hypoxic lung cancer cells. A possible explanation for this may be inhibition of acetylcholinesterase by hypoxia induced mir-132 [21 - 23] leading to accumulation of acetylcholine in these cells. Acetylcholine being an endogenous ligand of *nAChR-α7* can activate the downstream signaling cascades involved in the acquisition of a malignant phenotype characterized by an increased risk of metastasis and lower response rates to chemotherapy and/or radiotherapy similar to that in hypoxia [24, 25]. In hypoxic conditions, a significant dose dependent increase in HIF-1α was observed on treatment with Acetylcholine while a marked decrease in the expression of HIF-1α was observed in presence of *nAChR-α7* antagonist, α-BTX. Therefore, it appears that Acetylcholine and HIF-1α positively regulate each other. A noticeable increase in HIF-1α was also seen in the presence of *nAChR-α7* exogenous agonist nicotine in A549 lung cancer cells incubated in hypoxic condition affirming the response of the receptor to nicotine. Together these results suggest that *nAChR-α7* mediated signaling pathway and hypoxia inducible factors (HIFs), work together in hypoxic lung adenocarcinoma cells. The unchanged expression of HIF-2α in presence of both *nAChR-α7* antagonist and agonist together suggested that *nAChR-α7* mediated pathway regulates the expression of HIF-1α and not HIF-2α in these cells in hypoxia.

HIF-1α is well determined to be positively regulated via PI3K/AKT in many studies [26, 27]. In our study too, the decrease in expression of phosphorylated AKT in presence of α-BTX suggests that regulation of HIF-1α by *nAChR-α7* mediated pathway is through the PI3K/AKT pathway. Furthermore, blocking *nAChR-α7* mediated signaling pathway with its antagonist α-BTX inhibited hypoxia-induced migration in A549 cells thereby revealing a role for *nAChR-α7* in metastasis. Our findings confirm that hypoxia regulatory networks provide positive regulation that engages *nAchR* signaling, even in the absence of nicotine, to contribute to the disease prognosis and outcome.

This study puts forth a novel mechanism of hypoxia mediated increase in acetylcholine and *nAChR-α7* expression in lung cancer cells that positively regulates expression of HIF-1α via modulation of PI3K/AKT pathway and enhanced metastasis. A hypothetical model based on our results is shown in Figure 4. The study offers a new insight into the role of nAChR receptor signaling pathway independent of the smoking status and a plausible explanation of the similar disease progression in never-smokers and smokers. The insight gained from this study also opens up new avenues for drug development.

**Figure 4:**
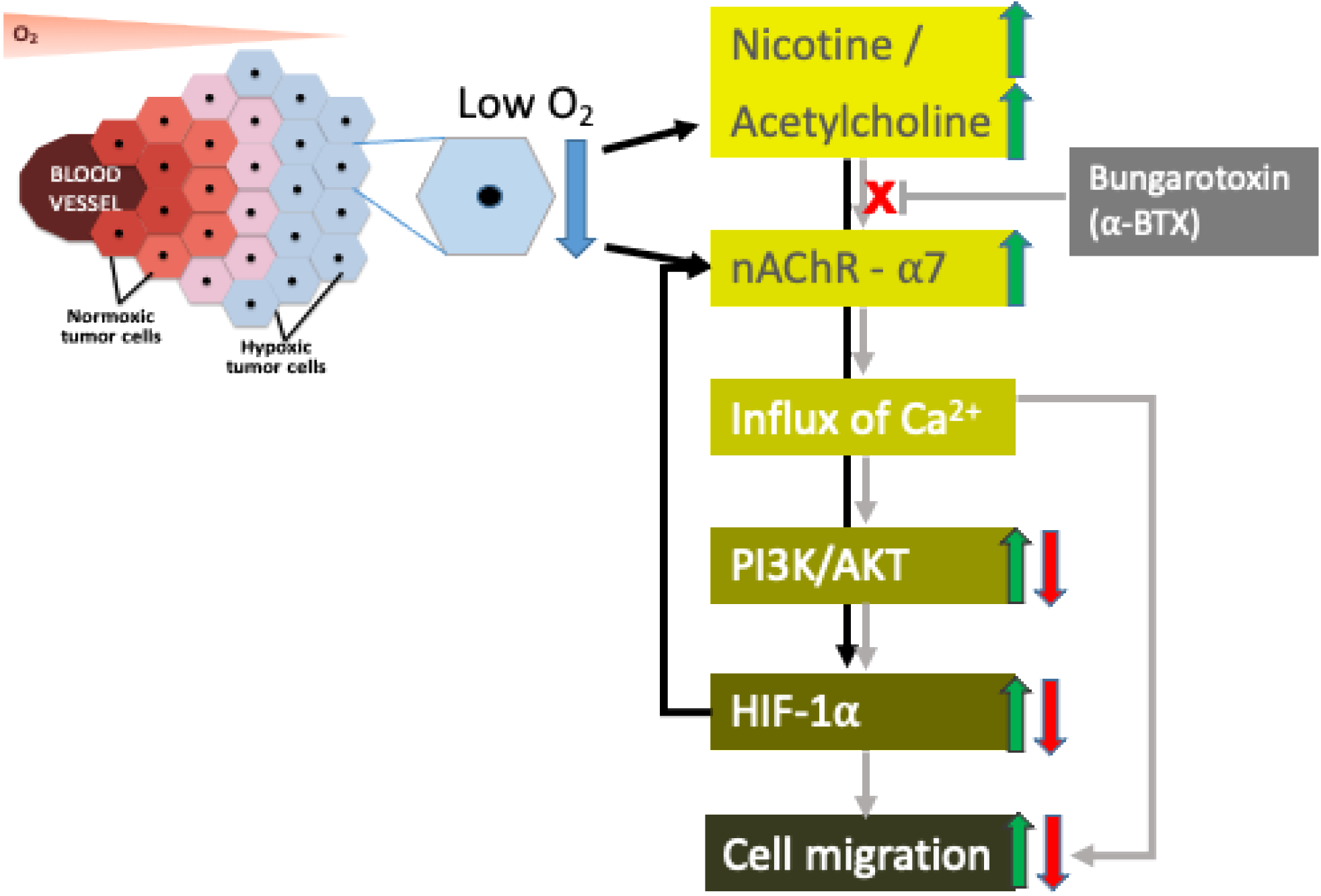
A model representing the proposed pathway by which nAChR-α7 mediated pathway and HIF-1α pathway interact with each other in A549 cells in hypoxia. Black arrows lines represent novel findings of our study. Grey arrows depict well established concepts. Green arrows show the upregulated pathways detected in this study. Inhibition by (α -BTX) and its downstream effect is represented with Red arrows.

## Acknowledgment

We thank Mr Shakeel Ansari for help in lab maintenance and Ms Kasthuri for help with maintaining lab accounts. NP was a recipient of UGC-BSR fellowship, JC is a recipient of DBT-JRF fellowship. Our lab is currently supported with grants from Science and Engineering Research Board and University funded Faculty Research Project, IOE scheme to TS. Funding support from DBT, ICMR, UGC for SAP (BRS III), DST for FIST (level II) program, DU-DST PURSE (phase II) is gratefully acknowledged. We thank Central Instrumentation Facility at University of Delhi South Campus for technical support.

